# Parallel patterns of cognitive aging in marmosets and macaques

**DOI:** 10.1101/2024.07.22.604411

**Authors:** Casey R. Vanderlip, Megan L. Jutras, Payton A. Asch, Stephanie Y. Zhu, Monica N. Lerma, Elizabeth A. Buffalo, Courtney Glavis-Bloom

## Abstract

As humans age, some experience cognitive impairment while others do not. When impairment does occur, it is not expressed uniformly across cognitive domains and varies in severity across individuals. Translationally relevant model systems are critical for understanding the neurobiological drivers of this variability, which is essential to uncovering the mechanisms underlying the brain’s susceptibility to the effects of aging. As such, non-human primates are particularly important due to shared behavioral, neuroanatomical, and age-related neuropathological features with humans. For many decades, macaque monkeys have served as the primary non-human primate model for studying the neurobiology of cognitive aging. More recently, the common marmoset has emerged as an advantageous model for this work due to its short lifespan that facilitates longitudinal studies. Despite their growing popularity as a model, whether marmosets exhibit patterns of age-related cognitive impairment comparable to those observed in macaques and humans remains unexplored. To address this major limitation for the development and evaluation of the marmoset as a model of cognitive aging, we directly compared working memory ability as a function of age in macaques and marmosets on the identical working memory task. Our results demonstrate that marmosets and macaques exhibit remarkably similar age-related working memory deficits, highlighting the value of the marmoset as a model for cognitive aging research within the neuroscience community.

## INTRODUCTION

Aging affects multiple cognitive domains in humans. While some individuals experience significant cognitive decline as they age, others maintain their cognitive abilities well into their later years (Small et al., 1999; Maher et al., 2022; Stern et al., 2023; Vanderlip et al., 2024). This variability highlights the importance of understanding the neurobiological mechanisms underlying age-related cognitive impairment. Identifying these mechanisms will be crucial for developing effective interventions and treatments for age-related cognitive impairment and age-related diseases, such as Alzheimer’s disease.

The use of animal models, particularly non-human primates, is critical in aging research due to their close genetic, physiological, and behavioral similarities to humans (Izpisua Belmonte et al., 2015). For decades, macaques have been the primary non-human primate model for studying cognitive aging (Gray and Barnes, 2019). Macaques have the ability to perform complex cognitive tasks and substantial work has employed the macaque model to understand the underlying neural mechanisms that support cognitive functions, including memory, attention and executive function. Further their patterns of age related cognitive decline closely mirror the impairments observed in humans and they spontaneously develop age-related neuropathologies such as neural hyperexcitability and increased deposition of beta amyloid and phosphorylated tau (Morrison and Baxter, 2012; Thomé et al., 2016; Arnsten et al., 2021). However, the long lifespan of the macaque poses logistical and practical challenges, particularly in conducting longitudinal studies that are essential for understanding the progression of cognitive impairment over time.

In this context, the common marmoset (*Callithrix jacchus*) has recently emerged as an alternative model for neuroscience research, particularly in the case of studies seeking to understand the biology of aging. Marmosets are the shortest lived anthropoid primate living 10 to 12 years and are considered aged at 7-8 years old. Therefore, they offer a pragmatic solution for longitudinal studies, which are less feasible in the longer-lived macaque, and are critically important for understanding aging as a biological process that unfolds over an extended timeframe (Tardif et al., 2011). Indeed, marmosets also exhibit age-related cognitive impairment and undergo age-related changes in the prefrontal cortex and hippocampus (Leuner et al., 2007; Glavis-Bloom et al., 2022, 2023). Yet, the suitability of marmosets as a model for age-related cognitive impairment remains underexplored. Critically, it is not well established whether marmosets exhibit patterns of cognitive decline with age that are comparable to those observed in macaques and humans. Studies have shown that marmosets have age-related impairments on tasks that are similar to those used to investigate cognitive impairment in macaques (Glavis-Bloom et al., 2022; Vanderlip et al., 2023). However, other studies have suggested that marmosets exhibit age-related cognitive decline in cognitive domains that are relatively unaffected by age in macaques (Rothwell et al., 2022).

Despite the growing popularity of the marmoset, comparative cognition studies with other non-human primate species are scarce (Nummela et al., 2019; Kell et al., 2023). Further, to-date, no study has directly investigated age-related cognitive impairment in marmosets and macaques performing the identical cognitive task. This work is critically needed to evaluate the extent to which marmosets and macaques exhibit similar cognitive repertoires. Therefore, our study investigated the similarities and differences in age-related cognitive impairment in the macaque and marmoset. To do this, we utilized the Delayed Recognition Span Task (DRST), which is a complex working memory task that requires the prefrontal cortex and hippocampus, two areas that are affected early in the aging process, and in Alzheimer’s disease (Beason-Held et al., 1999; Bor et al., 2006; Jeneson et al., 2010; Small et al., 2011). Previous work has demonstrated that older adults and macaques are impaired on this task compared to young controls, and people with Alzheimer’s disease are further impaired on this task compared to age-matched controls (Salmon et al., 1989; Herndon et al., 1997; Moss et al., 1997; Maylor et al., 2006; Belham et al., 2013; Mazurek et al., 2015; Satler et al., 2015). Further, we previously showed that marmosets could perform this complex task and that older marmosets were impaired across multiple aspects of the DRST (Glavis-Bloom et al., 2022). Utilizing this task, we conducted the first direct comparison of age-related cognitive impairment between marmosets and macaques; the two most commonly utilized primate species. This approach not only contributes to our understanding of cognitive aging in non-human primates, but also evaluates the potential of marmosets as a viable model for studying the neurobiology of age-related cognitive impairment. Comparative studies such as these are essential for advancing our understanding of cognitive decline mechanisms, ultimately guiding the development of targeted interventions and therapies for age-related cognitive disorders.

## METHODS

### Subjects

#### Marmosets

A total of 16 common marmosets (*Callithrix jacchus*) of both sexes participated in this study (8 female, 8 male). The marmosets ranged between 3.05 and 14.64 years of age at the onset of the study. Marmosets were housed singly or in pairs and were provided with species appropriate enrichment and diet. All procedures were carried out in accordance with the National Institutes of Health guidelines and were approved by the Salk Institute for Biological Studies Institutional Animal Care and Use Committee.

#### Macaques

A total of five female rhesus macaques (*Macaca mulatta*) participated in this study. Two of the monkeys were young (5.73 and 5.74 years of age), and three were aged (19.90, 20.70, and 23.66 years of age). All macaque monkeys were housed singly or in pairs in standard caging and were provided with species appropriate enrichment and diet. All procedures were carried out in accordance with the National Institutes of Health guidelines and were approved by the University of Washington Institutional Animal Care and Use Committee.

### Equipment

Cognitive testing for macaques and marmosets was administered via home cage mounted touch screen testing stations (Lafayette Instrument Company, Lafayette, IN). These stations were self-contained and included an infrared touch screen (15 inches, 764 x 1028 pixels for macaques; 10.4 inches, 800 x 600 pixels for marmosets) and reward delivery system (pellet dispenser for macaques; peristaltic pump for liquid rewards for marmosets). Cognitive tasks were programmed using Animal Behavior Environment Test (ABET) Cognition software (Lafayette Instrument Company, Lafayette, IN) that controlled all aspects of the task including the order of trials, timing, stimuli selection and display location, and delivery of rewards. The software also recorded detailed logs of task-related events (e.g., stimulus display, screen touches) with millisecond temporal resolution.

### Statistical analyses

Data were extracted from the ABET-produced logs and analyzed using custom purposed Python scripts. Non-parametric statistical analyses were used throughout the study. Spearman’s rank-order correlations were used to assess the relationship between age and various dependent variable performance metrics, and between task parameters and performance. Scheirer Ray Hare Tests were used to assess two factor interactions across time with Wilcoxon’s signed-rank tests or Mann Whitney U post-hoc tests. Friedman’s Tests with Nemenyi post-hoc tests were used to identify within-factor differences. Performance was compared to chance using Chi-Square Goodness of Fit Tests, and Mann Whitney U tests were used for species comparisons.

### Cognitive testing

#### Touch Training

All marmoset and macaque monkeys were naive to touch screen cognitive testing at the onset of the study. Therefore, all monkeys of both species were trained to operate the touch screens via a positive reinforcement procedure. Briefly, monkeys learned, through trial and error, that interacting with the touch screen yielded rewards. For marmosets, to encourage initial physical engagement with the screen, Marshmallow Fluff™ was applied in each of the nine locations where a blue square stimulus was presented. Once the monkeys associated touching the screen with earning rewards, no additional Marshmallow Fluff™ was applied (Glavis-Bloom et al., 2022). The macaques had previously been trained to touch a physical target to earn rewards. Therefore, to encourage initial physical engagement with the screen, the physical target was placed near the screen. Over the course of several days of training, the number of stimuli on the screen was reduced so that by the end of the touch training procedure, all monkeys were touching a single stimulus displayed on the screen in any of the possible locations. Then, over an additional few days of training, the amount of reward earned per screen touch was also reduced. Marmosets were rewarded with sweetened liquid such as apple juice, and macaques were rewarded with fruit-flavored pellets (190 mg Dustless Precision Pellets, Bio-Serv, Flemington, NJ).

#### Delayed Recognition Span Task (*Figure 1A*)

The Delayed Recognition Span Task (DRST) measures working memory capacity. Each trial of the DRST was initiated when a monkey touched a blue square in the center of the screen. Subsequently, a single black and white stimulus, chosen at random from a pool of 400 images, appeared on the screen in one of nine possible locations, also determined randomly (see Figure 1A for example stimuli). Upon touching this initial stimulus, the monkey received a small reward. After a delay, during which the screen remained blank, a two-alternative forced choice was presented. This choice included the original stimulus in its original location and a novel, visually distinct stimulus placed in a different pseudo-randomly selected location. If the monkey selected the novel stimulus, a correct response was recorded, a reward was dispensed, and another delay ensued. Subsequently, the first two stimuli reappeared in their original positions, and a third novel stimulus was introduced in a pseudo-randomly chosen location, with reward dispensed for selection of this new stimulus. This process continued with the introduction of novel stimuli after additional delays until the trial reached one of three possible conclusions: 1) the monkey successfully made nine consecutive correct selections; 2) the monkey failed to make a selection within a 12-second timeframe (i.e., omission); 3) the monkey made an incorrect response by selecting a non-novel stimulus. In cases of omission or incorrect responses, no reward was provided, and a five-second time-out period commenced before a new trial could be initiated. The "Final Span Length" for each trial was recorded as the number of correctly selected stimuli before the trial’s conclusion. The variations in the number of stimuli on the screen as trials progressed were referred to as trial difficulty levels (TDLs). Macaques and marmosets performed the DRST with a 2 second delay until performance levels plateaued. Subsequently, all macaques and a subset of the marmosets were tested on the DRST with delays greater than two seconds. Macaques and marmosets were tested with delays of 2, 6, 10, and 14 seconds, and macaques were additionally tested with a 30 second delay.

**Figure 1.**
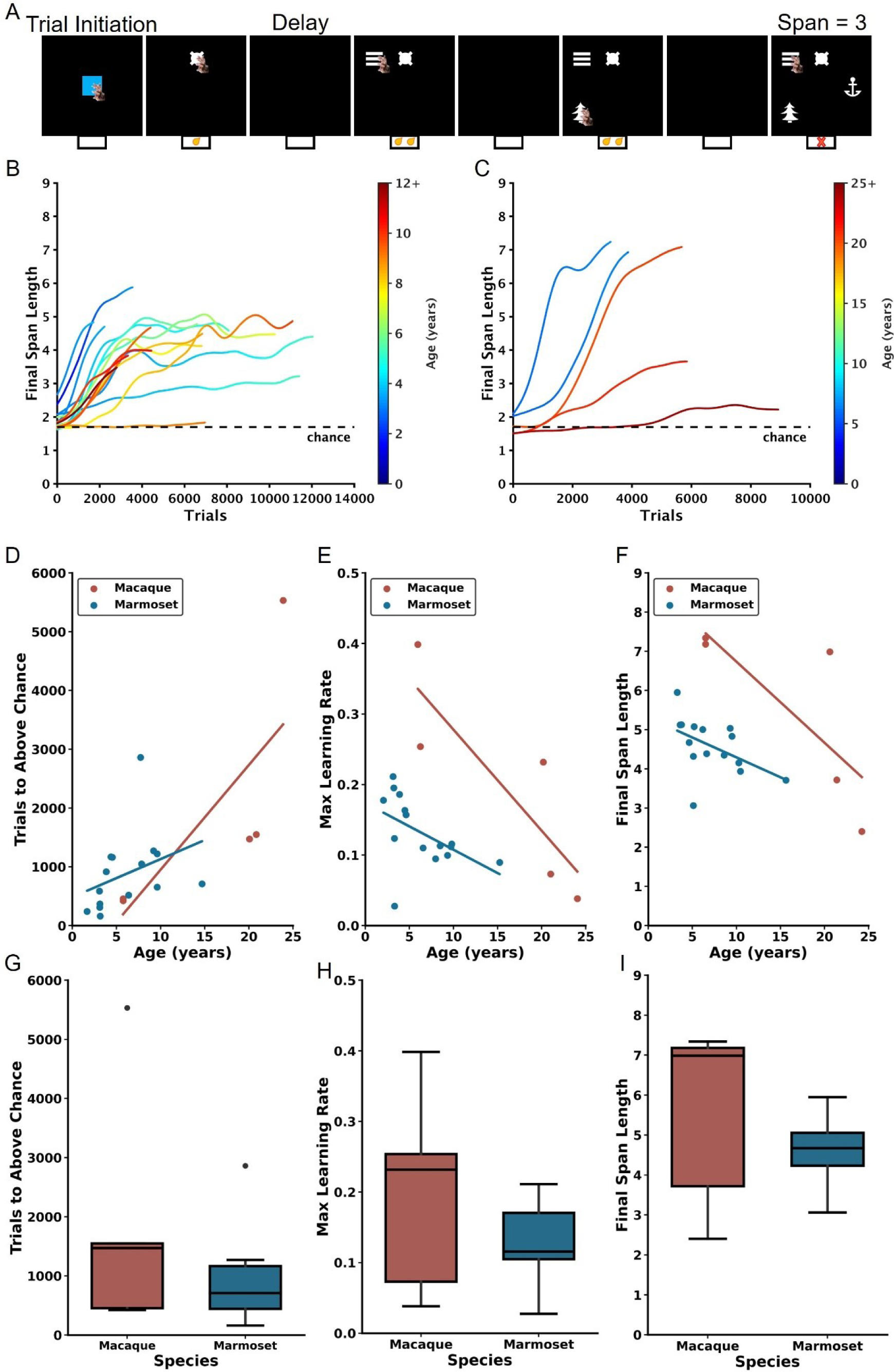
Age-dependent impairment in performance on the DRST in macaques and marmosets. A) Depiction of a single DRST trial. B) Marmoset and C) macaque individual learning curves. Each line denotes an individual animal, with color indicating the age during testing. The dashed black line represents chance level performance. Correlations show that increasing macaque (red) and marmoset (blue) age is associated with D) more trials needed to perform above chance in the Novice Phase, E) reduced maximum learning rates in the Learning Phase, and F) smaller working memory capacity. When averaging across ages, no interspecies differences were observed in G) trials to above-chance performance, H) maximum learning rates, or I) working memory capacity. Each circle in D-F represents one individual. Ages in D-F correspond to the age at the time of assessment. Black circles in boxplots in G-I represent outliers.

To maintain engagement and motivation, the quantity of reward increased in correspondence with the difficulty level of the trials. Specifically, marmosets received 0.05mL of reward for accurate responses when one stimulus was on the screen, 0.1mL for accurate responses when two, three, or four stimuli were on the screen, and 0.2mL for accurate responses when five, six, seven, eight, or nine stimuli were on the screen. Likewise, macaques earned one reward pellet when responding to one stimulus on the screen, earned two reward pellets for correct responses when two, three, or four stimuli were on the screen, and three reward pellets for correctly responding when five or more stimuli were on the screen.

Each marmoset and macaque underwent testing sessions two to five days per week, and each session concluded after three hours or once the marmoset had earned 20mL of reward, whichever event occurred first. Macaques underwent testing sessions three to five days a week and each session concluded after an hour or once the macaque earned 600 pellets, whichever event occurred first.

## RESULTS

Some of the marmoset data presented here are a subset of that published previously (Glavis-Bloom et al., 2022).

### Similar age-related learning and working memory impairment identified in macaques and marmosets

Consistent with previous studies employing the DRST (Herndon et al., 1997; Moss et al., 1997; Killiany et al., 2000; Moore et al., 2017), we used Final Span Length as the primary dependent measure for assessing performance. Final Span Length measures the working memory capabilities of the monkeys by recording the number of stimuli they correctly selected as novel in each trial. To analyze the learning progress of each animal, the Final Span Length data across time were represented in learning curves. These curves were refined using a Gaussian-weighted moving average applied across 2000 consecutive trials (Figure 1B-C).

Each marmoset and macaque learning curve was divided into three distinct Phases to facilitate the analyses described below. The Phases were determined by identifying two points on each curve. The first point identified when, during the course of learning, performance exceeded chance, and was determined by comparing the distribution of Final Span Lengths from a sliding block of 100 consecutive trials with a null distribution of Final Span Lengths derived from a Monte Carlo Simulation that approximated chance performance. The second point identified the time course of asymptotic performance and was determined by calculating the 90th percentile of Final Span Lengths achieved. The Novice Phase consisted of all trials up to and including the first point. The Learner Phase consisted of all trials between the two points, and the Expert Phase consisted of all trials after the second point. Each of the three Phases were determined to represent significantly different performance levels, thereby validating the method of dividing performance in this manner (Friedman’s test: Χ^2^ = 32, p = 1.13×10^-7^; Nemenyi post-hoc tests: Novice vs Learner p = 0.01, Novice vs Expert p = 0.001, Learner vs Expert p = 0.01). One marmoset never achieved DRST performance levels above chance and was excluded from these and all future analyses. Their learning curve is included in Figure 1B for illustrative purposes.

In the Novice Phase, monkeys began at chance levels of performance and gradually improved. There were significant positive associations between age and trials to above chance performance for both macaques and marmosets (Figure 1D; macaques: Spearman’s r(3) = 0.900, p = 3.739×10^-2^, marmosets: Spearman’s r(13) = 0.629, p = 8.988×10^-3^). Interestingly, there was no significant difference in performance between the species when collapsed across age (Figure 1G; Mann Whitney U = 51.00, p = 8.99×10^-3^).

In the Learner Phase, defined as trials from above chance performance until the 90th percentile, there were significant negative correlations between age and maximum learning rate (i.e., largest increase in Final Span Length over 100 trials) for each of the species (Figure 1E; macaques: Spearman’s r(3) = −1.000, p = 1.404×10^-24^, marmosets: Spearman’s r(13) = −0.643, p = 9.740×10^-3^). When data were collapsed across age, there was no significant species difference (Figure 1H; Mann Whitney U = 47, p = 0.44).

In the Expert Phase, where performance was between the 90th and 100th percentiles, for both macaques and marmosets there was a significant negative association between age and the maximum final span length achieved (Figure 1F; macaques: Spearman’s r(3) = −1.000, p = 1.404×10^-24^; marmosets: Spearman’s r(13) = −0.607, p = 1.638×10^-2^). When data were collapsed across age, there was no significant species difference (Figure 1I; Mann Whitney U = 47, p = 0.44).

### Associations between age and Delayed Non-Match-to-Sample performance

The first two parts of a DRST trial (Trial Difficulty Level (TDL)1 and TDL2, respectively) approximate a Delayed Non-Match-to-Sample (DNMS) paradigm. Specifically, the single stimulus presented in TDL1 is akin to a DNMS sample, and the two stimuli presented in TDL2 are akin to a DNMS choice. Therefore, by measuring performance of monkeys on DRST TDL2 trials, we can estimate DNMS task acquisition in the context of the DRST task. The two most frequently used dependent measures to assess DNMS performance are errors and trials to a learning criterion. We set the criterion a posteriori at 90% accuracy, achieved by responding correctly on at least 18 out of 20 consecutive trials. Spearman correlations revealed strong, significant associations between age and both errors to criterion (ETC) and trials to criterion (TTC) for both macaques and marmosets (Figure 2A, ETC; macaques: r(3) = 1.00, p = 1.40×10^-24^; marmosets: r(13) = 0.52, p = 4.78×10^-2^, Figure 2C, TTC; macaques: r(3) = 1.00, p = 1.40×10^-24^; marmosets: r(13) = 0.58, p = 2.37×10^-2^). Direct comparisons of macaque and marmoset DNMS performance revealed similar levels, whether measured by ETC or TTC, and regardless of age (Figure 2B, ETC: Mann Whitney U = 47.00, p = 0.44; Figure 2D, TTC: Mann Whitney U = 43.00, p = 0.67).

**Figure 2.**
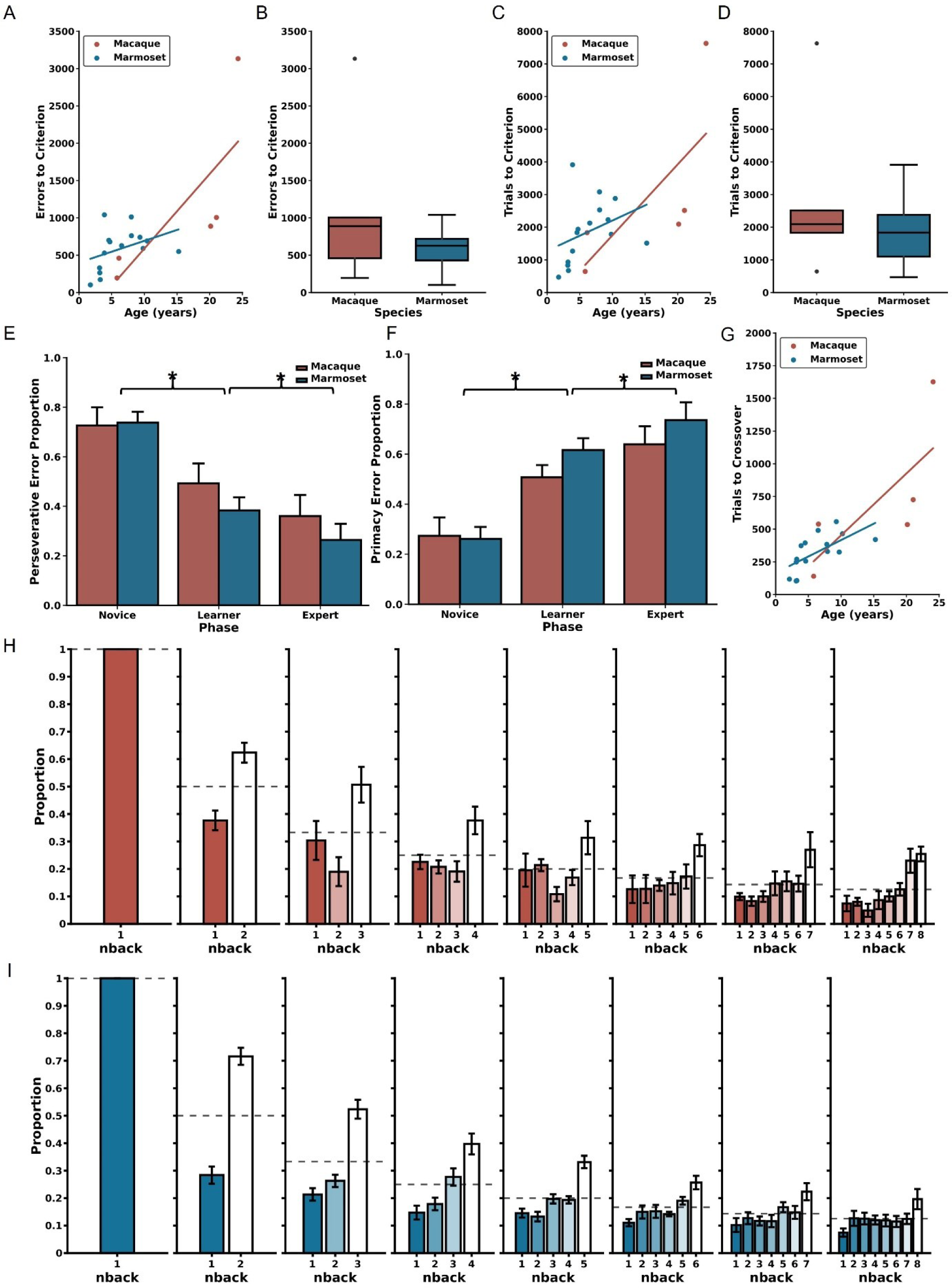
Error patterns related to age and trial difficulty level. A) Increased age is correlated with committing a larger number of errors before reaching the performance criterion in the DNMS section of the DRST for both macaques (red) and marmosets (blue). B) There were no significant species-specific differences in errors to reach the criterion. Similar patterns were identified in trials to criterion with C) increased age associated with requiring more trials needed to reach criterion. D) There were no significant interspecies differences in the number of trials to criterion. E) Reduction in perseverative errors across Novice, Learner, and Expert Phases in both species. F) Concurrent increase in primacy errors observed through these Phases for both macaques and marmosets. G) There were significant associations between increasing age and more trials to transition from predominantly perseverative to predominantly primacy errors for macaques (red) and marmosets (blue). H) During the Expert Phase, macaques more frequently misidentified remote (higher n-back) stimuli as novel compared to recent stimuli (lower n-back), suggesting retroactive interference. I) Marmosets exhibit a similar pattern during the Expert Phase, also suggesting vulnerability to retroactive interference; mean ± SEM, *p < 0.05.

### Similar age-related changes in error type process scores identified in macaques and marmosets

Trials ending with a Final Span Length of two present a unique opportunity to investigate the types of errors the monkeys made during each of the Phases of the DRST. This is because ending a trial with a Final Span Length of two entails that an error was made when there were three stimuli on the screen (i.e., TDL3). One of the stimuli is the correct choice, and the two other stimuli, when chosen, are each incorrect. If an error is made by choosing the stimulus that has just been rewarded on TDL2, this is a “perseverative” error, whereas if an error is made by choosing the stimulus that was rewarded earliest in the trial, on TDL1, this is a “primacy” error.

In marmosets, we previously reported that the proportion of perseverative errors decreased across the Phases of the DRST, whereas the proportion of primacy errors increased across the Phases (Glavis-Bloom et al., 2022). When assessing the performance of both marmosets and macaques for the proportion of perseverative errors, we found no significant main effect of Species, but there was a significant main effect of Phase, and a significant Species by Phase interaction (Figure 2E-F; Scheirer Ray Hare: Species H(1) = 0.86, p = 0.35; Phase H(2) = 43.71, p = 3.22×10^-10^, Species x Phase interaction H(2) = 45.47, p = 1.34×10^-10^). Similar results were found for proportion of primacy errors (Figure 2E-F; Scheirer Ray Hare: Species H(1) = 0.86, p = 0.35; Phase H(2) = 43.71, p = 3.22×10^-10^; Species x Phase interaction H(2) = 224.91, p = 1.45×10^-49^). The significant effects of Phase were driven by significant decreases in Perseverative errors across Phases, and corresponding increases in primacy errors across Phases (Wilcoxon signed-rank tests; Novice vs Learner p = 3.81×10^-6^; Learner vs Expert p = 0.001; Novice vs Expert p = 3.05×10^-5^). Together, these results suggest that while species alone did not significantly affect the proportion of perseverative or primacy errors, the Phase did, and the impact of Phase differed depending on the Species. Specifically, marmosets had a greater shift away from perseverative and towards primacy errors as they learned the task.

Overall, the switch from making predominantly perseverative to predominantly primacy errors occurred in the Learning Phase. To assess in more fine-grain detail the time course of this change in predominant error type, we measured the number of trials prior to the equivalence point where the monkeys made an equal proportion of the two error types. Doing so revealed strong and significant associations between age and trials to the equivalence point, for both species (Figure 2G; macaques: Spearman’s r(3) = 0.90, p = 3.74×10^-2^; marmosets: Spearman’s r(13) = 0.75, p = 1.39×10^-3^).

In the Expert Phase, trials of increased difficulty were completed consistently, providing the opportunity to investigate whether performance was affected by working memory interference. To explore this, we assessed how errors were distributed based on how far back in the trial’s history the incorrectly chosen stimulus was presented (referred to as "n-back"). For instance, if a monkey made a mistake when there were five objects on the screen (TDL5), they would receive a "Final Span Length" score of four for that trial. In this scenario, they could make an error by selecting the first object presented on the trial (n-back 4, known as primacy), the second object (n-back 3), the third object (n-back 2), or the fourth object (n-back 1, known as perseverative).

We quantified the distribution of n-back errors for each TDL and compared it to what would be expected by chance using Chi-Square Goodness of Fit Tests for marmosets and macaques separately (Figures 2H-I). The results of these tests demonstrated that, for all TDLs, the observed distributions of n-back errors significantly differed from what would be expected by chance for both macaques and marmosets (for statistics, see Table 1). Additionally, except for TDL3, there were no significant differences in the distribution of n-back errors made by macaques versus marmosets (for statistics, see Table 1). These findings indicate that both macaques and marmosets experienced retroactive interference, where newly acquired information disrupts the temporary storage of memories, resulting in errors when identifying stimuli presented earlier in the trial as if they were novel.

**Table 1.** Results from Chi-Square Goodness of Fit Tests to evaluate the distribution of n-back errors by TDL and species in the Expert Phase.

| <u>TDL</u> | <u>Macaque</u> |  | <u>Marmoset</u> |  | <u>Macaque vs Marmoset</u> |  |
| --- | --- | --- | --- | --- | --- | --- |
|  | <u>$\chi^2$</u> | <u>p-value</u> | <u>$\chi^2$</u> | <u>p-value</u> | <u>$\chi^2$</u> | <u>p-value</u> |
| 3 | 13.944 | $1.88 \times 10^{-4}$ | 340.682 | $4.533 \times 10^{-76}$ | 12.795 | $3.48 \times 10^{-4}$ |
| 4 | 25.064 | $3.609 \times 10^{-6}$ | 254.173 | $6.411 \times 10^{-56}$ | 3.164 | 0.206 |
| 5 | 10.680 | 0.0136 | 163.802 | $2.770 \times 10^{-35}$ | 4.211 | 0.240 |
| 6 | 16.118 | $2.87 \times 10^{-3}$ | 124.154 | $6.922 \times 10^{-26}$ | 5.604 | 0.231 |
| 7 | 19.757 | $1.39 \times 10^{-3}$ | 74.851 | $9.996 \times 10^{-15}$ | 3.026 | 0.696 |
| 8 | 15.727 | 0.0153 | 56.776 | $2.0278 \times 10^{-10}$ | 3.009 | 0.808 |
| 9 | 21.533 | $3.06 \times 10^{-3}$ | 30.166 | $8.854 \times 10^{-5}$ | 8.162 | 0.318 |

### Similar choice latency patterns between macaques and marmosets reveal effects of cognitive load

One of the benefits of using infrared touch screens for evaluating cognitive performance is their capability to precisely and consistently measure choice response times. This metric is widely recognized as a reliable indicator of processing speed, and it shows associations with cognitive load and task complexity (Romberg et al., 2013; Bopp and Verhaeghen, 2018; De Boeck and Jeon, 2019). To investigate whether this trend persisted when monkeys were engaged in the DRST, we analyzed the response times for correct and incorrect choices made by each monkey in various Phases of the DRST, as well as for different levels of task difficulty in the Expert Phase. A Scheirer Ray Hare Test uncovered significant main effects related to response type (correct choice, incorrect choice) and DRST Phase (Novice, Learner, Expert), along with a significant interaction between these factors (Figure 3A-B; response type: H(1)=22.635, p=1.959×10^-6^; phase: H(2)=14.954, p=5.66×10^-4^; interaction: H(2)=44.733, p=1.934×10^-10^). There was no main effect of species (H(1) = 0.159, p = 0.690).

**Figure 3.**
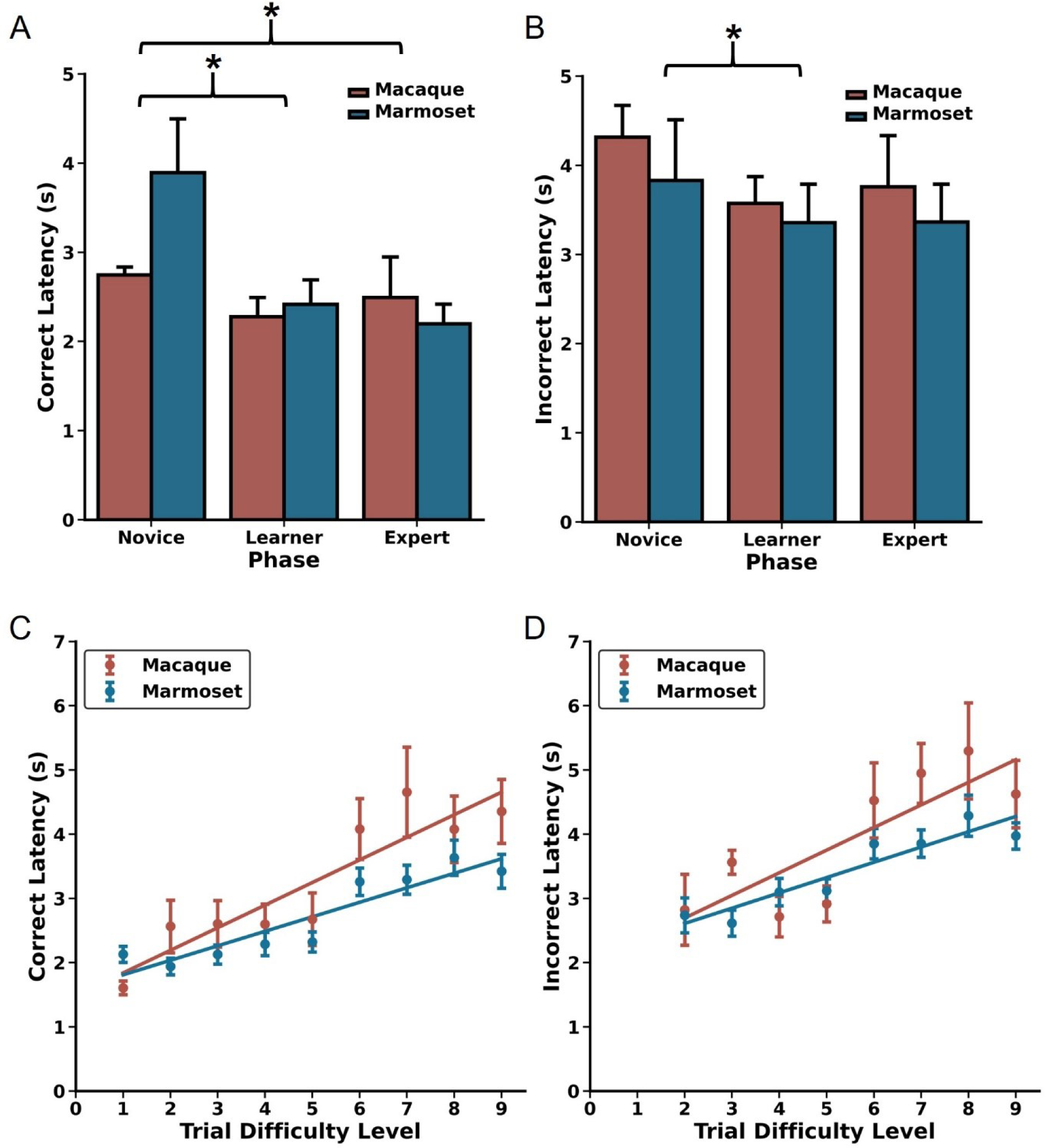
Choice latencies change as a function of DRST Phase and Trial Difficulty Level. Changes in A) correct choice latencies and B) incorrect choice latencies across the Novice, Learner, and Expert Phases for macaques (red) and marmosets (blue). Latencies decreased with increased task experience. In the Expert Phase, significant positive Spearman’s correlations were observed between trial difficulty level and C) correct choice latencies and D) incorrect choice latencies, reflecting increased cognitive load on more challenging portions of trials. mean ± SEM, *p < 0.05.

When investigating how correct choice latency changed as a function of experience, we found that, across species, correct latencies were longer during the Novice Phase compared to the Learner and Expert Phases, with no significant difference between the Learner and Expert Phases (Wilcoxon signed-rank test; Novice vs Learner: p = 3.0518×10^-5^, Novice vs Expert: p =5.80×10^-4^, Learner vs Expert: p = 0.117). We observed a similar pattern with incorrect latency, demonstrating that both macaques and marmosets make choices more rapidly as they gain experience and proficiency on the DRST (Wilcoxon signed-rank test; Novice vs Learner: p = 0.029, Novice vs Expert: p = 0.093, Learner vs Expert: p = 1.00). To explore whether incorrect choices might be attributed to impulsiveness, we compared the response times of correct and incorrect choices within each of the Phases. During the Novice Phase, correct and incorrect choice response times were similar. However, during the Learner and Expert Phases, incorrect choice response times were significantly longer than those for correct choices (Figure 3A-B; Wilcoxon signed-rank tests; Novice: p = 0.216; Learner: p = 1.907×10^-6^; Expert: p = 3.052×10^-5^). This suggests that when monkeys made errors, it was unlikely due to impulsivity, as they took considerably more time to respond in such instances.

We next examined whether elevated cognitive load was reflected in the choice latency data from the Expert Phase. To do this, we analyzed correct and incorrect choice latency data separately for each of the TDLs. We found strong, positive associations between increasing TDL and increasing correct latency for both macaques and marmosets (Spearman’s rank-order correlations; Macaque: r(7) = 0.900, p = 9.431×10^-4^; Marmoset: r(6) = 0.933, p = 2.359×10^-4^).

Similar associations were also found between TDL and incorrect latency (Spearman’s rank-order correlations; Macaque: r(6) = 0.810, p = 1.490×10^-2^; Marmoset: r(6) = 0.952, p = 2.604×10^-4^). Together, these results demonstrate that, as TDLs increase, so does cognitive load, and this is reflected in increased processing time and longer choice latencies in both macaques and marmosets.

### Species-specific effects of longer delays on DRST performance metrics

After reaching plateaued levels of performance on the DRST when trials included a 2 second delay between each stimulus presentation working memory was taxed further by the addition of longer delays. All macaques and a subset of the marmosets were tested with these longer delays which included 6, 10, and 14 seconds, and macaques were additionally tested with a 30 second delay.

First, we evaluated the effects of longer delays on DRST performance as measured by Final Span Length, and compared these effects across species. We found a significant main effect of species, no significant main effect of delay, and a significant species by delay interaction (Figure 4A; Scheirer Ray Hare: species H(1) = 19.25, p = 1.149×10^-5^; delay H(4) = 8.295, p = 0.0814, species x delay interaction H(4) = 70.189, p = 2.071×10^-14^). The significant main effect of species was driven by significant differences in performance between macaques and marmosets on all delays greater than 2 seconds (Mann-Whitney U tests; 2 seconds U = 18.0, p = 0.440, 6 seconds U = 5.0, p = 0.0127, 10 seconds U = 2.0, p = 6.21×10^-3^, 14 seconds U = 1.0, p = 8.66×10^-3^). These differences emerged because marmosets exhibited a significant delay-dependent decrease in Final Span Length, whereas macaque performance trended towards a delay-dependent decrease in Final Span Length but did not reach statistical significance (Wilcoxon signed-rank test results in Table 2).

**Figure 4.**
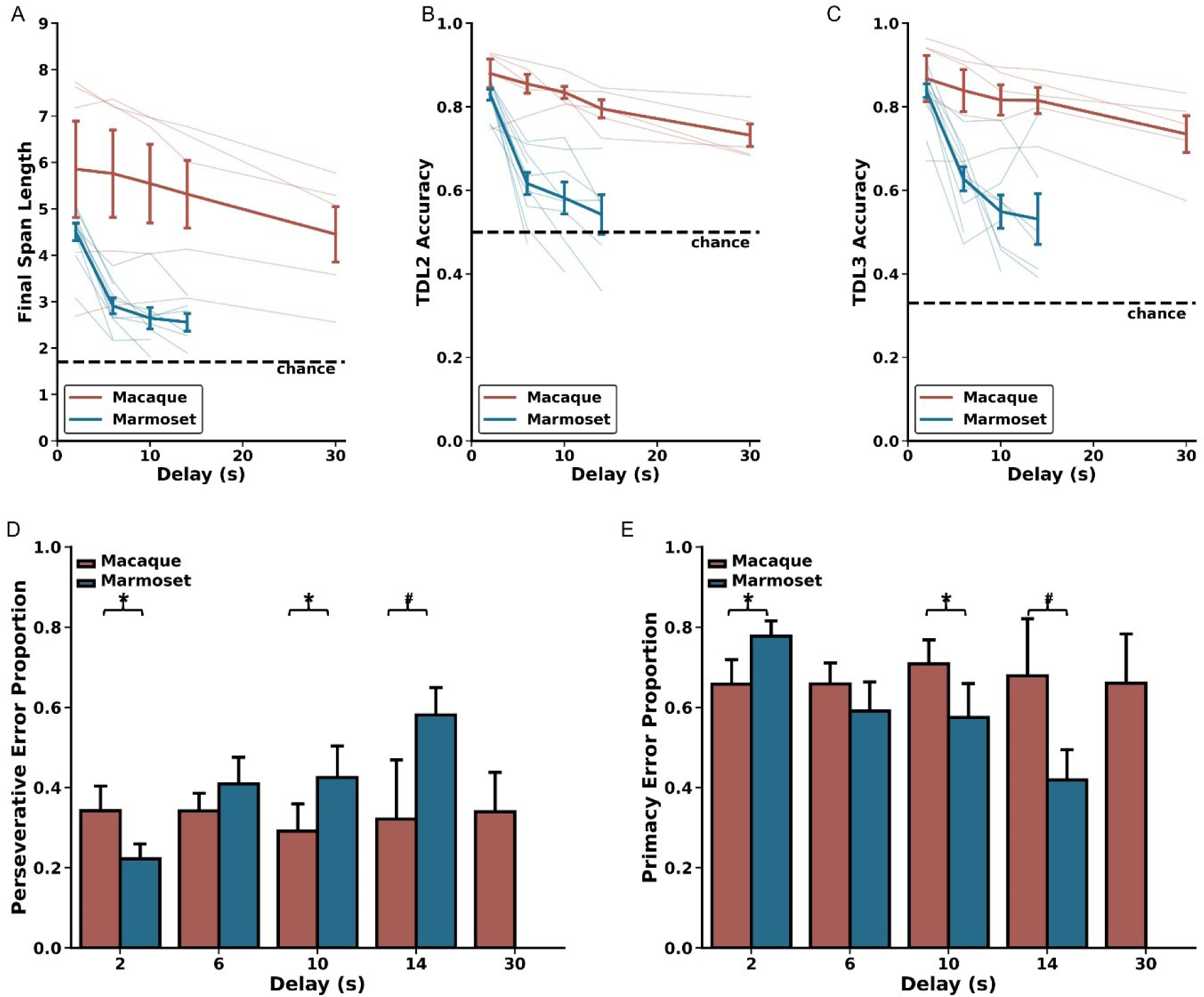
Delay-related effects on DRST performance. Marmosets (blue) show significant delay-dependent decreased DRST performance, whereas macaques (red) do not. Also, macaques have significantly higher performance than marmosets at delays longer than 2 seconds. These results are seen on several measures of performance including A) average Final Span Length, B) accuracy on the DNMS (TDL2) portion of the DRST, and C) accuracy on TDL3 trials. D) On TDL3 trials, marmosets’ perseverative errors increased in a delay-dependent manner, whereas macaque perseverative errors remained consistent across varying delays. E) Marmosets’ primacy error rate showed a corresponding delay-dependent decrease, and macaque primacy errors remained consistent across the varying delays. Lightly shaded lines in A-C depict individual animal performance as a function of delay. Bold colored lines in A-C depict species average performance as a function of delay. mean ± SEM, *p < 0.05.

**Table 2.** P-values for Wilcoxon signed-rank tests to compare performance across delays by species for Final Span Length (FSL), accuracy on Trial Difficulty Level (TDL)2 and TDL3, and proportion of perseverative errors.

| <u>Delay (s)</u> | <u>Macaques</u> |  |  |  | <u>Marmosets</u> |  |  |  |
| --- | --- | --- | --- | --- | --- | --- | --- | --- |
|  | <u>FSL</u> | <u>TDL2</u> | <u>TDL3</u> | <u>Errors</u> | <u>FSL</u> | <u>TDL2</u> | <u>TDL3</u> | <u>Errors</u> |
| 2 vs 6 | 0.813 | 0.0625 | 0.0625 | 1.0 | 0.00195 | 0.00195 | 0.00195 | 0.00195 |
| 2 vs 10 | 0.313 | 0.125 | 0.125 | 0.313 | 0.00781 | 0.00781 | 0.00781 | 0.0156 |
| 2 vs 14 | 0.313 | 0.188 | 0.188 | 1.0 | 0.0313 | 0.0313 | 0.0313 | 0.0313 |
| 2 vs 30 | 0.0625 | 0.0625 | 0.0625 | 1.0 | N/A | N/A | N/A | N/A |
| 6 vs 10 | 0.188 | 0.313 | 0.313 | 0.273 | 0.0781 | 0.0547 | 0.0547 | 1.0 |
| 6 vs 14 | 0.313 | 0.438 | 0.438 | 1.0 | 0.0625 | 0.0625 | 0.0625 | 0.156 |
| 6 vs 30 | 0.0625 | 0.0625 | 0.0625 | 1.0 | N/A | N/A | N/A | N/A |
| 10 vs 14 | 0.313 | 0.813 | 0.813 | 1.0 | 0.563 | 0.438 | 0.438 | 0.156 |
| 10 vs 30 | 0.0625 | 0.0625 | 0.0625 | 0.625 | N/A | N/A | N/A | N/A |
| 14 vs 30 | 0.0625 | 0.0625 | 0.0625 | 0.625 | N/A | N/A | N/A | N/A |

Next, we evaluated the effects of longer delays on TDL2 and TDL3 accuracy. We found significant main effects of species and delay and a significant species by delay interaction on both of these TDLs (TDL2: Figure 4B; Scheirer Ray Hare: species H(1) = 17.536, p = 2.819×10^-5^; delay H(4) = 17.449, p = 1.581×10^-3^; species x delay interaction H(4) = 110.182, p = 6.656×10^-23^; TDL3: Figure 4C; Scheirer Ray Hare: species H(1) = 14.239, p = 1.610×10^-4^; delay H(4) = 14.749, p = 5.250×10^-3^; species x delay interaction H(4) = 101.587, p = 4.518×10^-21^). The significant main effects of species were driven by significant differences in performance between macaques and marmosets on all delays greater than 2 seconds (Mann-Whitney U tests; TDL2: 2 seconds U = 10.0, p = 0.0753, 6 seconds U = 0.0, p = 6.660×10^-4^, 10 seconds U = 0.0, p = 1.554×10^-3^, 14 seconds U = 0.0, p = 4.329×10^-3^; TDL3: 2 seconds U = 17.0, p = 0.371, 6 seconds U = 4.0, p = 7.992×10^-3^, 10 seconds U = 2.0, p = 6.216×10^-3^, 14 seconds U = 1.0, p = 8.658×10^-3^). These differences emerged because marmosets exhibited a delay-dependent decrease in accuracy on both TDL2 and TDL3, whereas macaques did not (Wilcoxon signed-rank test results in Table 2). Together, these results suggest that the effect of delay on performance varied as a function of species.

As described above, trials ending with a Final Span Length of two present a unique opportunity to investigate the prevalence with which monkeys committed perseverative and primacy errors. We assessed the proportion of these types of errors as a function of species and delay length and found no main effect of species, no main effect of delay, but a significant species by delay interaction (Figure 4D-E; Scheirer Ray Hare: species H(1) = 1.329, p = 0.249; delay H(4) = 16.204, p = 0.0940; species x delay interaction H(4) = 309.889, p = 1.260×10^-60^). This significant interaction is driven by the fact that marmosets exhibited a delay-dependent increase in perseverative errors and a corresponding delay-dependent decrease in primacy errors, whereas macaque perseverative and primacy errors were unchanged across varied delays (Wilcoxon signed-rank test results in Table 2). Thus, the proportion of error types changed as a function of delay only in marmosets.

## DISCUSSION

In this study, we conducted the first direct comparison of cognitive ability as a function of age in macaques and marmosets. By testing young and aged macaques and marmosets on the identical working memory task, we found that they exhibit remarkably similar age-related learning and working memory impairments. This work establishes that the patterns of age-related working memory deficits are largely conserved across the two most common nonhuman primate models used for cognitive aging research. Macaques demonstrate more robust performance than marmosets when working memory is taxed through increased delay durations.

### Evaluation of macaque and marmoset performance in the context of prior work

In humans and non-human primates, cognitive functions that rely on the prefrontal cortex and hippocampus decline with age. As such, working memory deficits appear particularly early in the aging process (Gazzaley et al., 2005; Upright and Baxter, 2021). To measure working memory as a function of aging across macaques and marmosets, we used a touch screen version of the DRST. Using this task, we found strikingly similar associations between advancing age and working memory impairment in macaques and marmosets.

Specifically we found that aged macaques have impaired ability to acquire the rules of the DRST, requiring more experience to perform above the levels expected by chance, and learning at a slower rate, than young macaques. This parallels findings from previous investigations in the marmoset that demonstrated age-related impairments in acquisition and learning of the DRST (Glavis-Bloom et al., 2022). We also found age-related decreased working memory capacity in both macaques and marmosets. These findings align with previous work that has documented age-related impairments on the DRST in each of these species independently, albeit on similar, but non-identical task designs (Moss et al., 1997, 1997; Shobin et al., 2017; Glavis-Bloom et al., 2022; Moore et al., 2023).

We capitalized on the fact that within the context of each DRST trial there existed an opportunity to directly compare performance between macaques and marmosets on the more commonly-used DNMS paradigm. We found that, in both species, aging was associated with impaired performance, measured by errors to a learning criterion. This aligns with numerous studies in macaques reporting similar findings (Rapp and Amaral, 1989; Hara et al., 2012; Comrie et al., 2018; Baxter et al., 2023; Gray et al., 2023).

Although we find clear and compelling evidence for age-related working memory impairment, evaluation of individual animal learning curves revealed striking levels of between animal variability which was particularly evident in older individuals. Similar to previous reports in humans, similarly aged macaques and similarly aged marmosets demonstrated different working memory aptitudes. A subset of animals of each species performed at high levels, while others performed less optimally.

Given that a subset of the macaques approached ceiling levels of performance when 2 second delays were employed, we evaluated potential differences in DRST performance between macaques and marmosets under more challenging experimental conditions. Historically, working memory performance decreases as a function of longer delays that tax working memory (Beason-Held et al., 1999; Dumitriu et al., 2010; Comrie et al., 2018; Baxter et al., 2023). We found that increasing delays affected the performance of marmosets and macaques differently. Whereas macaques maintained stable levels of performance on delays up to 30 seconds, marmoset performance was severely impaired by increased delays. Future work is needed to understand the neural mechanisms that explain this species difference.

### Process scores reveal similar cognitive mechanisms underlying age-related impairment in macaques and marmosets

Process scores refer to metrics that provide insight into the cognitive processes underlying performance on a task, beyond the final outcome score (Kaplan, 1988). Process scores are valuable because, in humans, they can predict future cognitive decline (Thomas et al., 2018; Edmonds et al., 2019). Previously, we demonstrated that process scores are critical to revealing the specific mechanisms that contribute to age-related cognitive impairments in marmosets (Glavis-Bloom et al., 2023; Vanderlip et al., 2023). Here, we used process scores to determine whether macaques and marmosets showed working memory impairment due to shared underlying cognitive mechanisms.

The types of errors (perseverative vs primacy) committed while performing a working memory task are indicative of the strategy used to perform the task. Early in the learning process, we found that monkeys predominantly made perseverative errors. This likely results from application of a “win-stay” strategy prior to an understanding of the DRST rules that necessitate “win-shift” to correctly choose a novel object. We found that with increased experience and performance on the DRST, both macaques and marmosets shifted from making predominantly perseverative errors to predominantly primacy errors. Further, in both species, older age is associated with a protracted shift between predominant error types. This aligns with prior work showing that both aged macaques and marmosets take longer to shift from a "win-stay" strategy to a "win-shift" strategy on other cognitive tasks (Moore, 2003; Sadoun et al., 2019). Finally, we found that once monkeys had enough task experience to perform at high levels on the DRST and were making predominantly primacy errors, they did so by selecting stimuli encountered earliest in the trial sequence. This pattern shows that macaques and marmosets both succumb to retroactive interference, indicating that the errors made are not random, but rather reflect specific cognitive interference processes underlying performance metrics.

Processing speed is associated with cognitive load and task complexity and can be measured via choice latencies (Bopp and Verhaeghen, 2018; De Boeck and Jeon, 2019). The use of infrared touch screen systems in our study facilitated reliable capture of choice latencies with 1ms temporal resolution, enabling us to evaluate any potential species differences reflected in this process score (Bussey et al., 2008; Smith et al., 2023). We found, in both species, that choice latencies were longer when the response was incorrect than when the response was correct. These findings demonstrate that incorrect responses were not a result of impulsivity, and therefore support the idea that the age-related impairments on performance metrics reflect valid measurement of cognitive ability. Further, we found that choice latencies increased as a function of increased task difficulty in both macaques and marmosets. This affirms that macaques and marmosets experience similarly increased cognitive loads across trial difficulty levels on the DRST.

### Underlying biological mechanisms of age-related working memory impairment

Substantial research in macaques has revealed age-related alterations in the dorsolateral prefrontal cortex (dlPFC) that may underlie working memory impairment. In particular, age-related synapse loss in the dlPFC is associated with impaired working memory (Peters et al., 2008; Dumitriu et al., 2010). Moreover, this synapse loss is driven by a specific decrease in the number of small synapses which critically support working memory (Dumitriu et al., 2010; Arnsten et al., 2012; Dickstein et al., 2013). Additionally, dysmorphic changes in synaptic mitochondria within the dlPFC are also associated with impairments in working memory in aging macaques (Hara et al., 2014). In contrast, research on age-related changes in the marmoset dlPFC is relatively limited. However, our previous findings (Glavis-Bloom et al., 2023) have shown that aged marmosets, similar to macaques, exhibit synapse loss, which is predominantly due to a decrease in small synapses. Further, we discovered that age-related impairments on the DRST were linked to a mismatch in the sizes of synaptic mitochondria and their corresponding boutons in aged marmosets. This mismatch, or lack of coordination, is believed to cause a decoupling effect, leading to an imbalance between energy supply and demand, and ultimately resulting in impaired synaptic transmission and working memory impairment (Glavis-Bloom et al., 2023).

Significant age-related changes that are correlated with memory impairment is also evident in the hippocampus of aged macaques. Unlike the dlPFC, the macaque hippocampus does not exhibit an overt age-related loss of synapses (Hara et al., 2012). There are, however, age-related changes in the number of synapses per bouton in the dentate gyrus, and in aged macaques, an increase in non-synaptic boutons correlates with recognition memory impairment (Hara et al., 2011). Strikingly, there has been extremely limited investigation of age-related changes in the marmoset hippocampus. Although there are a few studies that report reduced neurogenesis and increased phosphorylated tau (Leuner et al., 2007; Rodríguez-Callejas et al., 2016; Perez-Cruz and Rodriguez-Callejas, 2023), there are no studies linking age-related hippocampal changes to cognitive function.

Our study provides the first direct comparison of age-related cognitive impairments between macaques and marmosets, revealing that these species exhibit similar learning and working memory deficits with age. The observation that macaque working memory performance is more resilient to the effects of longer delays suggests a potentially larger working memory capacity compared to marmosets. Future work is needed to understand whether similar neural circuits underlie performance on this task across these species, and also to determine what age-related neuropathology gives rise to declining working memory.

## Acknowledgments

This research was supported by an AHA-Allen Initiative in Brain Health and Cognitive Impairment award made jointly through the American Heart Association and The Paul G. Allen Frontiers Group: 19PABH134610000AHA, National Institutes of Health grants 1R21AG068967-01 and P51OD010425, grants from the Larry L. Hillblom Foundation and the Don and Lorraine Freeberg Foundation, and the Fiona and Sanjay Jha Chair in Neuroscience. We thank Katie Williams for assistance in the care of the marmosets and technical support. We thank Kelly Morrisroe for assistance in training of the rhesus macaques and Paige Robertson for assistance with behavioral task programming.

